# YAP/TAZ mediate TGFβ2-induced Schlemm’s canal cell dysfunction

**DOI:** 10.1101/2022.06.06.494681

**Authors:** Haiyan Li, Ayushi Singh, Kristin M. Perkumas, W. Daniel Stamer, Preethi S. Ganapathy, Samuel Herberg

## Abstract

**Purpose:** Elevated transforming growth factor beta2 (TGFβ2) levels in the aqueous humor have been linked to glaucomatous outflow tissue dysfunction. Potential mediators of dysfunction are the transcriptional co-activators, Yes-associated protein (YAP) and transcriptional coactivator with PDZ binding motif (TAZ). However, the molecular underpinnings of YAP/TAZ modulation in Schlemm’s Canal (SC) cells under glaucomatous conditions are not well understood. Here, we investigate how TGFβ2 regulates YAP/TAZ activity in human SC (HSC) cells using biomimetic extracellular matrix (ECM) hydrogels, and examine whether pharmacologic YAP/TAZ inhibition would attenuate TGFβ2-induced HSC cell dysfunction.

**Methods:** Primary HSC cells were seeded atop photocrosslinked ECM hydrogels, made of collagen type I, elastin-like polypeptide and hyaluronic acid, or encapsulated within the hydrogels. HSC cells were induced with TGFβ2 in the absence or presence of concurrent actin destabilization or pharmacologic YAP/TAZ inhibition. Changes in actin cytoskeletal organization, YAP/TAZ activity, ECM production, phospho-myosin light chain levels, and hydrogel contraction were assessed.

**Results:** TGFβ2 significantly increased YAP/TAZ nuclear localization in HSC cells, which was prevented by either filamentous (F)-actin relaxation or depolymerization. Pharmacologic YAP/TAZ inhibition using verteporfin without light stimulation decreased fibronectin expression and reduced actomyosin cytoskeletal rearrangement in HSC cells induced by TGFβ2. Similarly, verteporfin significantly attenuated TGFβ2-induced HSC cell-encapsulated hydrogel contraction.

**Conclusions:** Our data provide evidence for a pathologic role of aberrant YAP/TAZ signaling in HSC cells under simulated glaucomatous conditions, and suggest that pharmacologic YAP/TAZ inhibition has promising potential to improve outflow tissue dysfunction.

## Introduction

The Schlemm’s canal (SC) is a continuous vessel that encircles the anterior chamber at the iridocorneal angle; its lumen is lined with a single non-fenestrated layer of endothelial cells having both blood and lymphatic characteristics ^1–5^. Situated in close apposition to the trabecular meshwork (TM), the SC is divided into the inner and outer wall ^2^. The SC inner wall experiences a basal-to-apical pressure gradient (intraocular versus episcleral venous pressures) that drives aqueous humor into the SC lumen, which is then drained into the collector channels and aqueous veins ^6^. Most of the resistance to aqueous humor outflow is generated at, or close to the SC inner wall in a region called the juxtacanalicular tissue (JCT) of the TM ^7–9^. Importantly, increased outflow resistance in the JCT leads to elevated intraocular pressure (IOP), the only modifiable risk factor for primary open-angle glaucoma (POAG) ^8,10–13^.

Previous studies have demonstrated that glaucomatous SC cells isolated from POAG eyes exhibited higher levels of filamentous (F)-actin, α-smooth muscle actin (αSMA), and fibronectin, as well as increased cell stiffness compared to normal SC cells ^14,15^. The ocular hypertension-causing steroid dexamethasone was shown to increase F-actin fibers, while the IOP-lowering Rho-associated kinase (ROCK) inhibitor Y27632 decreased F-actin levels ^16,17^; F-actin is thought to mediate SC cell contractility and stiffness to negatively affect aqueous humor outflow and IOP ^18^. Thus, SC cell dysfunction is thought to be a significant contributor to the increased outflow resistance in POAG; however, the mechanistic underpinnings of SC cell pathobiology remain incompletely understood.

Transforming growth factor beta 2 (TGFβ2), the predominant TGFβ isoform in the eye and aqueous humor, is a major player in contributing to the pathologic changes in POAG ^10,19–23^. It has been shown that levels of TGFβ2 are elevated in eyes of glaucoma patients compared to age-matched normal eyes ^21,22,24,25^. In culture, TM and SC cells isolated from donor eyes with POAG secrete more active TGFβ2 protein compared to normal cells ^15,26^. Accordingly, perfusion of TGFβ2 in human anterior segments increases resistance in the conventional outflow pathway ^27^. At the cellular level, TGFβ2 increases actin stress fibers and phospho-myosin light chain (p-MLC) to drive pathologic TM cell contraction ^28–30^. Moreover, exposure of TM cells to TGFβ2 induces the expression/deposition of extracellular matrix (ECM) proteins such as collagen types I and IV, and fibronectin ^28,31–33^. Despite the progress made in uncovering the role of TGFβ2 in TM cell dysfunction, the contributions of TGFβ2 to SC cell pathobiology are less well understood.

Yes-associated protein (YAP) and transcriptional coactivator with PDZ-binding motif (TAZ) are powerful regulators of cell proliferation and differentiation, with an established link to tissue fibrosis ^34–38^. Upon nuclear translocation, YAP/TAZ interact with TEA domain (TEAD) transcription factors to drive the expression of known (CTGF, CYR61, ANKRD1) and glaucoma-related putative downstream effectors of active YAP/TAZ signaling (e.g., transglutaminase-2 (TGM2)) ^39^. To that end, *YAP1* was recently identified among a group of previously unknown POAG risk loci across European, Asian, and African ancestries, suggesting a potential causal association with outflow dysfunction ^40^. YAP/TAZ can be activated by multiple stimuli, such as stiffened ECM, increased mechanical stress, and exposure to growth factors ^41,42^. Recently, we demonstrated that both ECM stiffening and TGFβ2 increase YAP/TAZ activity in human TM cells, which was linked to pathologic cell contractility and ECM remodeling that may exacerbate glaucoma pathology ^26^.

The physiological substrate of the SC inner wall endothelial cells is its discontinuous basal lamina, which is in direct contact with the JCT of the TM ^3,6^. Recently, we developed a bioengineered hydrogel composed of ECM biopolymers found in the native JCT region encapsulated with human TM (HTM) cells, and demonstrated its utility for studying cell-ECM interactions under normal and simulated glaucomatous conditions in a relevant tissue-mimetic 3D microenvironment ^26,30,43^. Here, we investigated the effects of TGFβ2 on regulating YAP/TAZ activity in primary human SC cells cultured atop the acellular ECM hydrogel. Additionally, we examined whether pharmacologic YAP/TAZ inhibition would alleviate TGFβ2-induced cell dysfunction.

## Materials and Methods

### HSC cell isolation and culture

Experiments using human donor eye tissue were approved by the SUNY Upstate Medical University Institutional Review Board (protocol #1211036), and were performed in accordance with the tenets of the Declaration of Helsinki for the use of human tissue. Primary human SC (HSC) cells were isolated from healthy donor corneal rims discarded after transplant surgery and cultured according to established protocols ^5^. Briefly, the corneal rims were cut into wedges and placed into a 100 mm dish containing low-glucose Dulbecco’s Modified Eagle’s Medium (DMEM; Gibco; Thermo Fisher Scientific, Waltham, MA, USA) with 10% fetal bovine serum (FBS; Atlanta Biologicals, Flowery Branch, GA, USA) and 1% penicillin/streptomycin/glutamine (PSG; Gibco). Using an SMZ1270 stereomicroscope (Nikon Instruments, Melville, NY, USA), a 2% gelatin-coated (Sigma-Aldrich) 6-0 nylon monofilament sterile suture (eSutures, Mokena, IL, USA) was inserted into the SC lumen of each wedge with fine-tipped forceps (Fine Science Tools, Foster City, CA, USA), and these wedges were cultured in DMEM with 10% FBS and 1% PSG, and maintained at 37°C in a humidified atmosphere with 5% CO_2_ for 3 weeks. Next, curvilinear incisions were made parallel to Schwalbe’s line (alongside the suture) into the TM, which produced a TM flap. After lifting the TM, the sutures were gently removed, and a ~2 mm long piece was cut from each end of the suture to prevent fibrotic cell contamination. Next, the sutures were washed with Dulbecco’s Phosphate Buffered Saline (DPBS; Gibco), placed into a single well of gelatin-coated (Sigma-Aldrich) 6-well culture plates (Corning; Thermo Fisher Scientific), and digested for 2 min with 0.25% trypsin/0.5 mM EDTA (Gibco). Subsequently, 5 ml DMEM with 10% FBS and 1% PSG were added, and the sutures were moved to another gelatin-coated well. The digests and sutures were cultured in DMEM with 10% FBS and 1% PSG, and maintained at 37°C in a humidified atmosphere with 5% CO_2_. Fresh media was supplied every 2-3 days. Once confluent, HSC cells were lifted with 0.25% trypsin/0.5 mM EDTA and sub-cultured in DMEM with 10% FBS and 1% PSG. All studies were conducted using cells passage 3-6. Four HSC cell strains (HSC01, HSC02, HSC03, HSC09) were characterized and used for the experiments herein; the reference cell strain HSC78 was isolated and characterized at Duke University by K.M.P. and W.D.S. (**Table 1**). Different combinations of 3 HSC cell strains were used per experiment, depending on cell availability.

**Table 1:** HSC cell strain information.

| ID | Sex | Age |
| --- | --- | --- |
| HSC01 | Male | 33 |
| HSC02 | Male | 46 |
| HSC03 | Female | 46 |
| HSC09 | Female | 69 |
| Reference HSC78* | Male | 77 |
| Reference HSC88* | Female | 67 |
| *\*obtained from W.D.S. at Duke University* |  |  |

### HSC cell characterization

HSC cells were seeded at 1 × 10^4^ cells/cm^2^ in 6-well culture plates or on sterilized glass coverslips in 24-well culture plates, and cultured in DMEM with 10% FBS and 1% PSG. HSC cell morphology and growth characteristics were monitored by phase contrast microscopy using an LMI-3000 Series Routine Inverted Microscope (Laxco; Thermo Fisher Scientific). Once confluent, monolayers of HSC cells were processed for immunocytochemistry and immunoblot analyses to assess expression of fibulin-2 and vascular endothelial-cadherin, respectively, with human umbilical vein endothelial cells (ATCC, Manassas, VA, USA) serving as a positive control and previously validated HTM cells (HTM01 or HTM19) ^43^ serving as negative controls. To rule out the contamination of HTM cells, dexamethasone (Fisher Scientific)-induced myocilin expression was assessed in HSC cells. Briefly, confluent HSC cells were treated with 100 nM dexamethasone or vehicle control (0.1% (v/v) ethanol) in DMEM with 1% FBS and 1%PSG for 4 d, and serum- and phenol red-free DMEM for 3 d. The HSC cell culture supernatants were collected and concentrated using Amicon^®^ Ultra Centrifugal Filters (Millipore Sigma, Burlington, MA, USA) for immunoblot analysis.

### Preparation of ECM thin-film hydrogels

Hydrogel precursors methacrylate-conjugated bovine collagen type I (MA-COL, Advanced BioMatrix, Carlsbad, CA, USA; 3.6 mg/ml [all final concentrations]), thiol-conjugated hyaluronic acid (SH-HA, Glycosil^®^, Advanced BioMatrix; 0.5 mg/ml, 0.025% (w/v) photoinitiator Irgacure^®^ 2959; Sigma-Aldrich, St. Louis, MO, USA) and in-house expressed elastin-like polypeptide (SH-ELP, thiol via K<u>C</u>TS flanks ^43^; 2.5 mg/ml) were thoroughly mixed. Thirty microliters of the hydrogel solution were pipetted onto a Surfasil (Fisher Scientific)-coated 18 × 18-mm square glass coverslip followed by placing a regular 12-mm round glass coverslip onto the hydrogels. Constructs were crosslinked by exposure to UV light (OmniCure S1500 UV Spot Curing System; Excelitas Technologies, Mississauga, Ontario, Canada) at 320-500 nm, 2.2 W/cm^2^ for 5 s, as previously described ^26,30,43^. The hydrogel-adhered coverslips were removed with fine-tipped tweezers and placed in 24-well culture plates (Corning; Thermo Fisher Scientific). The elastic modulus of the ECM hydrogel was previously measured at ~0.2 kPa ^26^.

### HSC cell treatments

HSC cells were seeded at 2 × 10^4^ cells/cm^2^ atop premade ECM hydrogels and cultured in DMEM with 10% FBS and 1% PSG for 1 or 2 days. Then, HSC cells were cultured in serum-free DMEM with 1% PSG and subjected to the different treatments: TGFβ2 (2.5 ng/ml; R&D Systems, Minneapolis, MN, USA; for 3 d), the ROCK inhibitor Y27632 (10 μM; Sigma-Aldrich; for 3 d or 30 min only), the actin depolymerizer latrunculin B (10 μM; for 30 min only to preserve cell viability; Tocris Bioscience; Thermo Fisher Scientific), or the YAP inhibitor verteporfin (0.5 μM; Sigma-Aldrich; for 3 d).

### Immunoblot analysis

Equal protein amounts (10 or 40 μg), determined by standard bicinchoninic acid assay (Pierce; Thermo Fisher Scientific), from concentrated HSC cell culture supernatants ± dexamethasone at 7 d or from HSC cell lysates in lysis buffer (radioimmunoprecipitation assay (RIPA) buffer: 150 mM sodium chloride, 1.0% Triton™ X-100, 0.5% sodium deoxycholate, 0.1% sodium dodecyl sulfate, 50 mM Tris, pH 8.0; all from Thermo Fisher Scientific) supplemented with Halt™ protease/phosphatase inhibitor cocktail (Thermo Fisher Scientific) in 4× loading buffer (Invitrogen; Thermo Fisher Scientific) with 5% beta-mercaptoethanol (Fisher Scientific) were boiled for 5 min, subjected to SDS-PAGE using NuPAGE™ 4-12% Bis-Tris Gels (Invitrogen; Thermo Fisher Scientific) at 120V for 80 min, and transferred to 0.45 μm polyvinylidene difluoride (Sigma; Thermo Fisher Scientific) or 0.2 μm nitrocellulose membranes (Bio-Rad, Hercules, CA, USA). Membranes were blocked with 5% bovine serum albumin (Thermo Fisher Scientific) in tris-buffered saline with 0.2% Tween®20 (Thermo Fisher Scientific), and probed with primary antibodies followed by incubation with HRP-conjugated or fluorescent secondary antibodies (LI-COR, Lincoln, NE, USA). Bound antibodies were visualized with the enhanced chemiluminescent detection system (Pierce) on autoradiography film (Thermo Fisher Scientific) or Odyssey® CLx imager (LI-COR). A list of all antibodies and their working dilutions can be found in **Supplementary Table 1**.

### Immunocytochemistry analysis

HSC cells atop ECM hydrogels subjected to the different treatments for 3 d were fixed with 4% paraformaldehyde (Thermo Fisher Scientific) at room temperature for 20 min, permeabilized with 0.5% Triton™ X-100 (Thermo Fisher Scientific), blocked with blocking buffer (BioGeneX), and incubated with primary antibodies, followed by incubation with fluorescent secondary antibodies; nuclei were counterstained with 4’,6’-diamidino-2-phenylindole (DAPI; Abcam). Similarly, cells were stained with Phalloidin-iFluor 488 (Invitrogen) or 594 (Abcam)/DAPI according to the manufacturer’s instructions. Coverslips were mounted with ProLong™ Gold Antifade (Invitrogen) on Superfrost™ microscope slides (Fisher Scientific), and fluorescent images were acquired with an Eclipse N*i* microscope (Nikon Instruments, Melville, NY, USA). A list of all antibodies and their working dilutions can be found in **Supplementary Table 1**.

### Image analysis

All image analyses were performed using FIJI software (National Institutes of Health (NIH), Bethesda, MD, USA). Briefly, the cytoplasmic YAP/TAZ intensity was measured by subtracting the spatially overlapping nuclear (DAPI) intensity from the total YAP/TAZ intensity. The nuclear YAP/TAZ intensity was recorded as the proportion of total YAP/TAZ intensity that spatially overlapped with the nucleus (DAPI). YAP/TAZ nuclear/cytoplasmic (N/C) ratio was calculated as follows: N/C ratio = (nuclear YAP/TAZ signal/area of nucleus)/(cytoplasmic signal/area of cytoplasm). Fluorescence intensity of F-actin, FN, TGM2, and p-MLC were measured in at least 30 images from 3 HSC cell strains with 3 replicates per HSC cell strain with standardized image background subtraction (using species-matched IgG negative controls) using FIJI software. Given the lack of defined αSMA fibers in untreated controls, we quantified the number of cells exhibiting αSMA signal/fibers (i.e., yes/no decision) per image after standardized background correction and calculated % of αSMA-positive cells according to a previous report ^44^. At least 150 cells were analyzed in 30 images from 3 HSC cell strains with 3 replicates per HSC cell strain. For each experiment, care was taken to include an equivalent number of cells across groups.

### HSC hydrogel contraction analysis

HSC cell-laden hydrogels were prepared by mixing HSC cells (1.0 × 10^6^ cells/ml) with MA-COL (3.6 mg/ml), SH-HA (0.5 mg/ml, 0.025% (w/v) photoinitiator) and SH-ELP (2.5 mg/ml) on ice, followed by pipetting 10 μl droplets of the HSC cell-laden hydrogel precursor solution onto polydimethylsiloxane (PDMS; Sylgard 184; Dow Corning)-coated 24-well culture plates. Constructs were crosslinked as described above (320-500 nm, 2.2 W/cm^2^, 5 s). HSC cell-laden hydrogels were cultured in DMEM with 10% FBS and 1% PSG in presence of the different treatments. Longitudinal brightfield images were acquired at 0 d and 5 d with an Eclipse T*i* microscope (Nikon). Construct area from N = 12 hydrogels per group from 3 HSC cell strains with 4 replicates per cell strain was measured using FIJI software and normalized to 0 d followed by normalization to controls.

### HSC hydrogel cell viability analysis

Cell viability was measured with the CellTiter 96® Aqueous Non-Radioactive Cell Proliferation Assay (Promega) following the manufacturer’s protocol. HSC hydrogels cultured in DMEM with 10% FBS and 1% PSG in presence of the different treatments for 5 d were incubated with the staining solution (38 μl MTS, 2 μl PMS solution, 200 μl DMEM) at 37°C for 1.5 h. Absorbance at 490 nm was recorded using a spectrophotometer plate reader (BioTek, Winooski, VT, USA). Blank-subtracted absorbance values served as a direct measure of HSC cell viability from N = 12 hydrogels per group from 3 HSC cell strains with 4 replicates per cell strain.

### Statistical analysis

Individual sample sizes are specified in each figure caption. Comparisons between groups were assessed by two-way analysis of variance (ANOVA) with Tukey’s multiple comparisons *post hoc* tests, as appropriate. The significance level was set at p<0.05 or lower. GraphPad Prism software v9.2 (GraphPad Software, La Jolla, CA, USA) was used for all analyses.

## Results

### HSC cell characterization

Four HSC cell strains (HSC01, HSC02, HSC03, and HSC09) were used and compared to validated reference strains (HSC78 and HSC88). All of our HSC cell strains exhibited typical spindle-like elongated cell morphology comparable to the reference standard (**Fig. 1A**). A reliable feature of HSC cells *in vitro* is expression of two positive markers, fibulin-2 and vascular endothelial-cadherin (VE-Cad) ^45^. Our results showed that all HSC cell strains robustly expressed fibulin-2 (**Fig. 1B**) and VE-Cad (**Fig. 1C**), comparable to reference HSC cells, with HTM controls being negative for both markers ^45^. In culture, HTM cells upregulate myocilin expression following challenge with the corticosteroid dexamethasone ^46^, whereas this does not occur in HSC cells. We observed that none of the HSC cell strains expressed myocilin regardless of the presence or absence of dexamethasone (**Fig. 1D**), suggesting pure HSC cell preparations devoid of HTM cell contamination.

**Fig. 1.**
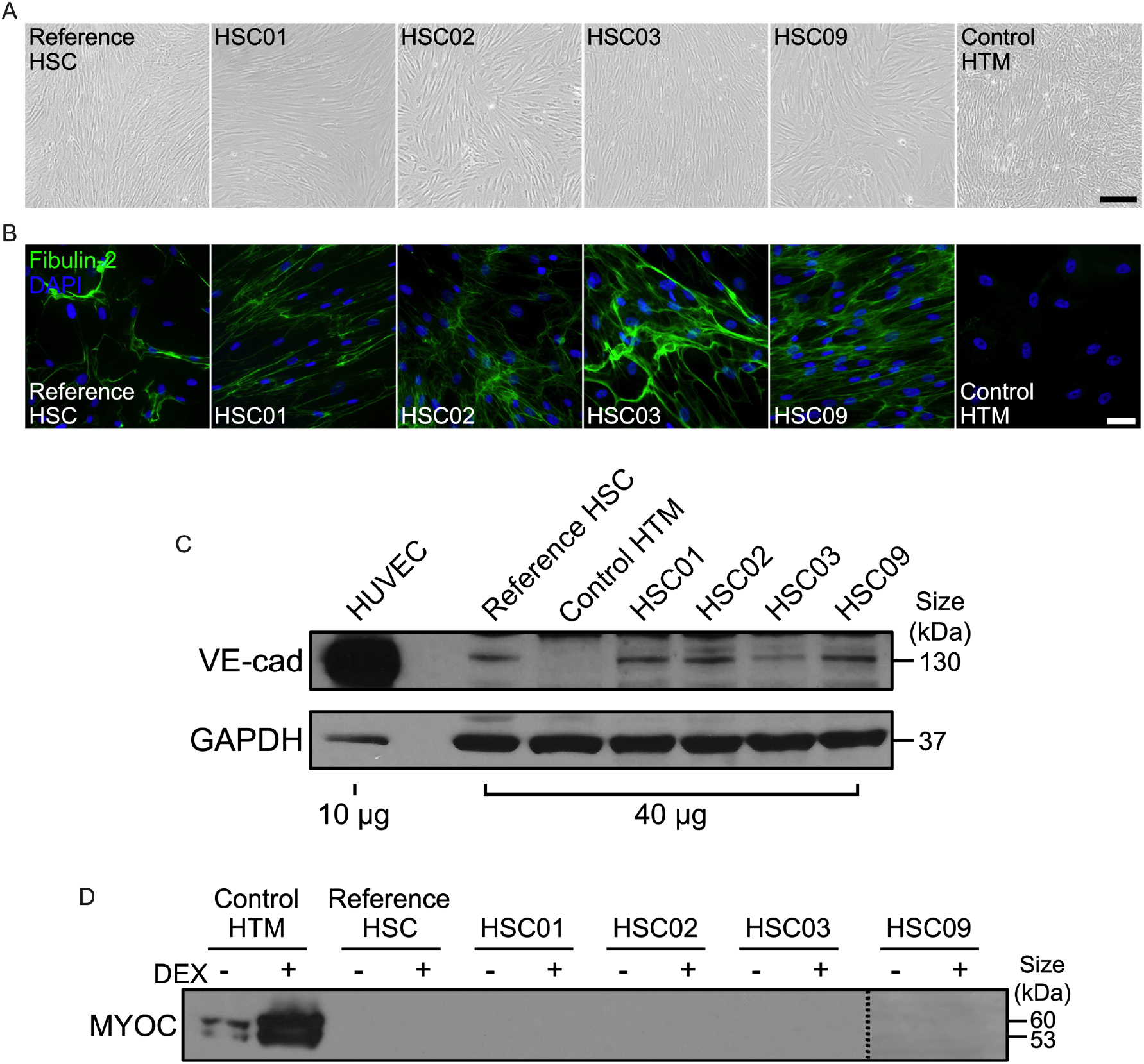
HSC cell characterization. (A) Representative phase contrast micrographs of reference HSC, HSC01, HSC02, HSC03 and HSC09 cell strains, with HTM controls. Scale bar, 200 μm. (B) Representative immunofluorescence micrographs of fibulin-2. Scale bar, 100 μm. (C) Immunoblot of VE-Cadherin (VE-Cad), with human umbilical vein endothelial (HUVEC) cells serving as positive control. (D) Immunoblot of secreted myocilin (MYOC) without or with dexamethasone (DEX) stimulation at 7 d (= negative marker). Reference HSC cells were provided by K.M.P./W.D.S.. Control HTM cells were previously characterized in ref ^43^.

Together, these data suggest that HSC01, HSC02, HSC03, and HSC09 exhibit required key characteristics according to previous publications ^5,45^ to faithfully identify them as normal HSC cells, comparable to confirmed reference standards.

### Effects of TGFβ2 in the absence or presence of ROCK inhibition or Lat B on actin cytoskeleton in HSC cells

F-actin stress fibers are found in higher numbers in SC cells isolated from POAG eyes compared to normal SC cells ^15^. Moreover, it has been shown that glaucomatous SC cells occasionally exhibit disorganized or “tangled” F-actin fibers; these abnormal cytoskeletal structures often appear as dense groups of actin bundles with multidirectional orientation and overlaps ^47^. To investigate the effects of TGFβ2 on actin cytoskeleton remodeling, HSC cells were cultured atop premade ECM hydrogels and treated with TGFβ2 alone or co-treated with ROCK inhibitor Y27632 or latrunculin B (Lat B), a compound that inhibits actin polymerization (**Fig. 2A**). We observed significantly increased F-actin stress fibers/overall intensity and qualitatively more disorganized F-actin filaments in HSC cells treated with TGFβ2 compared to controls, which was significantly prevented by co-treatment with Y27632 or Lat B, with short Lat B treatment for only 30 min showing a stronger effect (**Fig. 2B,C**). Of note, both types of actin destabilizers failed to fully block TGFβ2-induced F-actin assembly. We observed a range of responsiveness to the TGFβ2 challenge ± treatments among the HSC cell strains, indicating normal donor-to-donor viability.

**Fig. 2.**
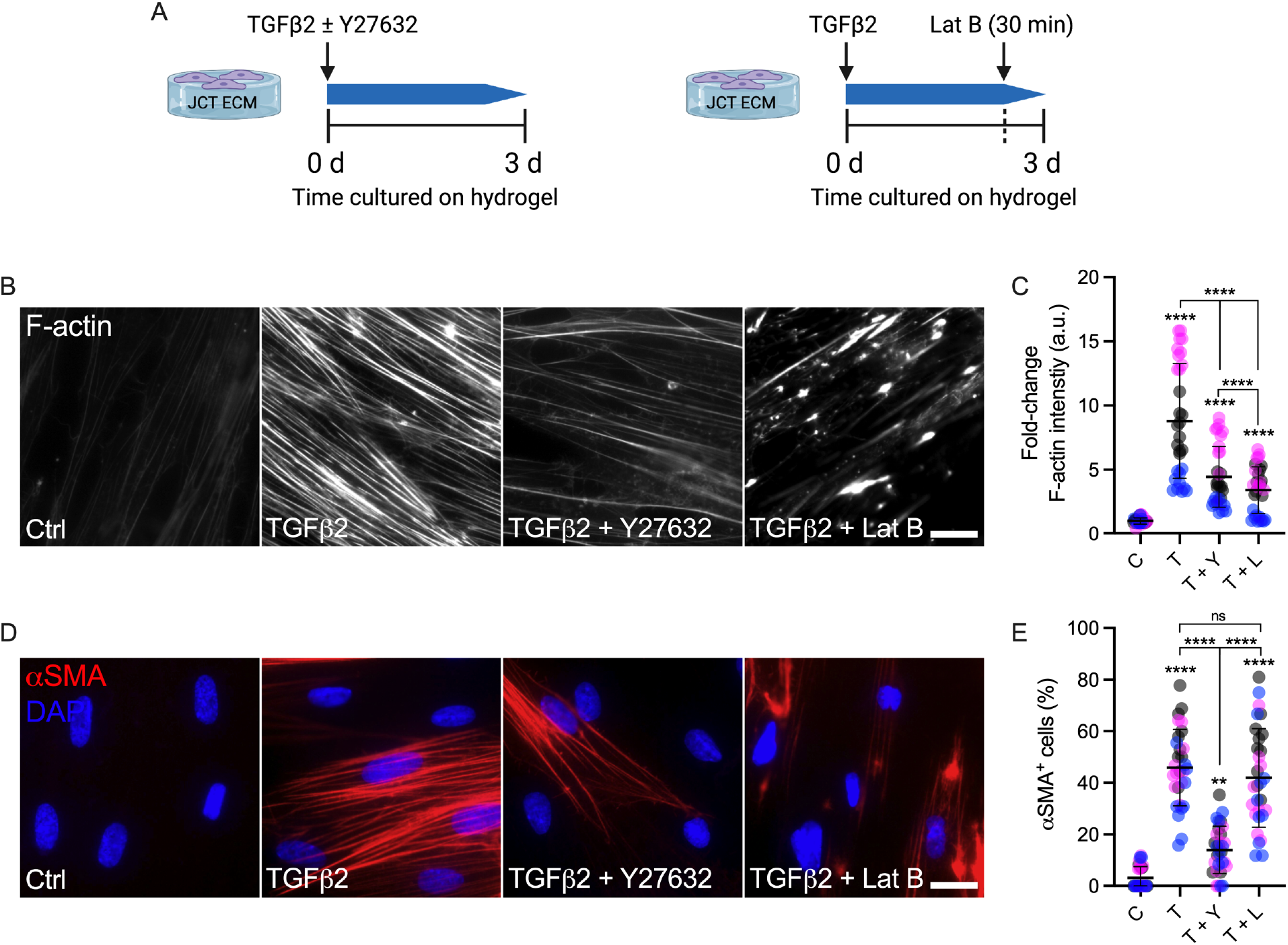
Effects of TGFβ2 in the absence or presence of a ROCK inhibitor or Lat B on F-actin and αSMA stress fibers in HSC cells. (A) Schematic showing time course of the different treatments. (B) Representative fluorescence micrographs of F-actin in HSC cells on ECM hydrogel substrates subjected to control, TGFβ2 (3 d; 2.5 ng/mL), TGFβ2 + Y27632 (3 d; 10 μM), TGFβ2 (3 d) + Lat B (30 min; 2 μM). Scale bar, 20 μm. (C) Analysis of F-actin intensity (N = 30 images per group from 3 HSC cell strains with 3 replicates per HSC cell strain). (D) Representative immunofluorescence micrographs of αSMA in HSC cells on ECM hydrogel substrates subjected to the different treatments. Scale bar, 20 μm. (E) Analysis of percentage of αSMA^+^ cells (N = 30 images per group from 3 HSC cell strains with 3 experimental replicates per HSC cell strain; more than 150 cells were analyzed per cell strain). Symbols with different colors represent different cell strains. The bars and error bars indicate Mean ± SD. Significance was determined by two-way ANOVA using multiple comparisons tests (**p<0.01, ****p < 0.0001. ns = non-significant).

We, and others, have shown that the fibrotic marker α-smooth muscle actin (αSMA) is upregulated in HTM cells by TGFβ2 exposure ^28–30,48^. Here, we demonstrated that TGFβ2 induced expression of αSMA in 45.87% of HSC cells compared to only ~3.16% in controls. Similar to results with F-actin, Y27632 prevented αSMA stress fiber formation induced by TGFβ2, with only 13.98%αSMA^+^ cells. Interestingly, short-term Lat B treatment failed to block TGFβ2-induced αSMA expression (% of αSMA^+^ cells: 41.95%) (**Fig. 2D,E**).

To ascertain that the results seen with concurrent exposure to TGFβ2 + Y27632 were not affected by waning inhibitor activity over the 3-day period, HSC cells were treated with TGFβ2 for 3 d followed by supplementation of Y27632 for only 30 min (**Fig. 3A**) to match the Lat B treatment regimen. We observed that HSC cells undergoing short-term ROCK inhibition with Y27632 fully blocked TGFβ2-induced F-actin stress fiber formation, restoring control levels (**Fig. 3B,C**). As such, ROCK inhibition for only 30 min appeared comparatively more potent than 3 d co-treatment (**Fig. 2B,C**). We also found that short-term supplementation of Y27632 significantly decreased expression of αSMA induced by TGFβ2 (**Fig. 3D,E**), consistent with the 3 d co-treatment data (**Fig. 2B,C**); yet, intensity levels remained significantly higher than controls.

**Fig. 3.**
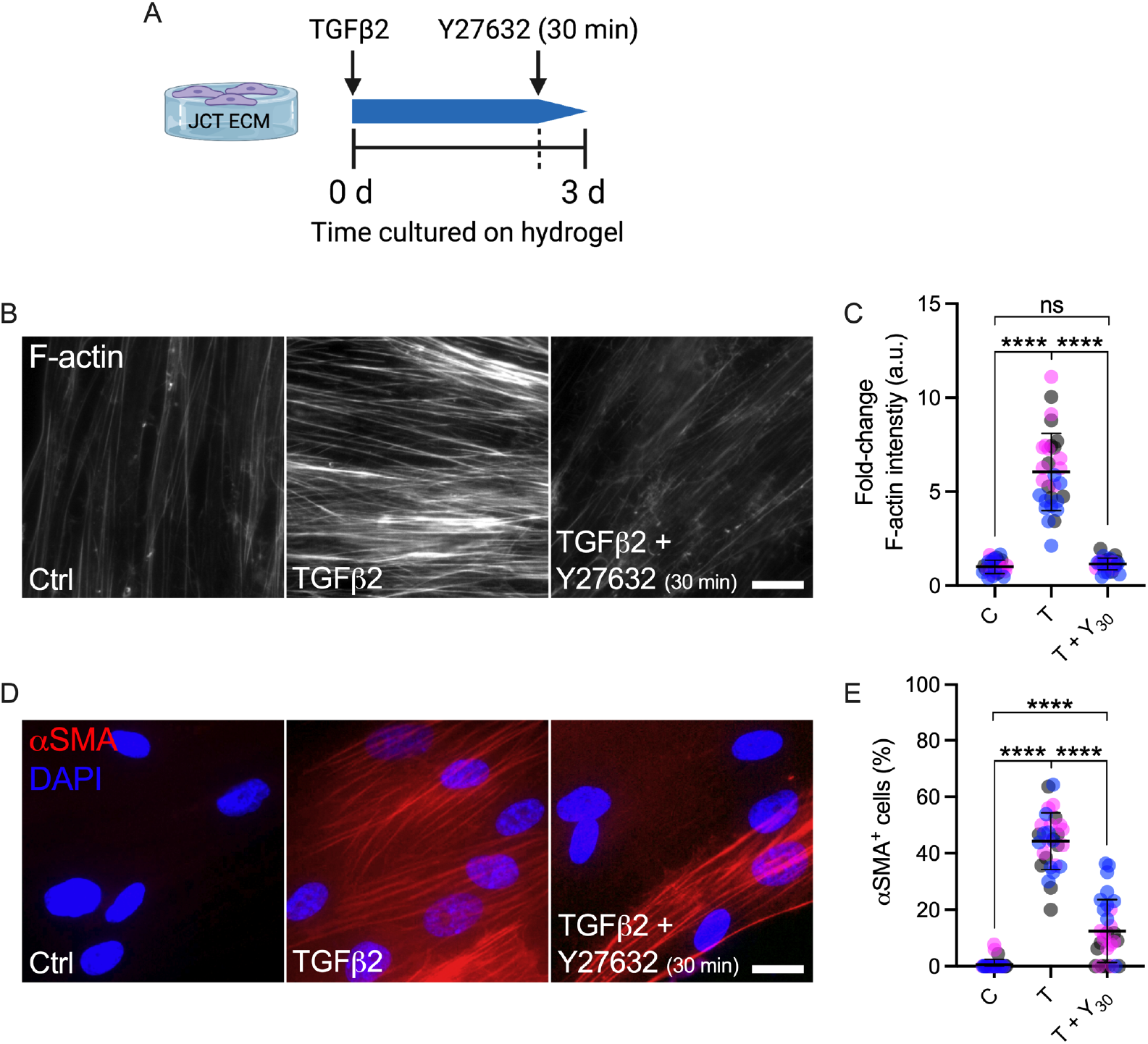
Effects of short-term ROCK inhibition on TGFβ2-induced F-actin and αSMA stress fibers in HSC cells. (A) Schematic showing time course of short-term ROCK inhibition. (B) Representative fluorescence micrographs of F-actin in HSC cells on ECM hydrogel substrates subjected to control, TGFβ2 (3 d; 2.5 ng/mL), TGFβ2 (3 d) + Y27632 (30 min; 10 μM). Scale bar, 20 μm. (C) Analysis of F-actin intensity (N = 30 images per group from 3 HSC cell strains with 3 replicates per HSC cell strain). (D) Representative immunofluorescence micrographs of αSMA in HSC cells on ECM hydrogel substrates subjected to the different treatments. Scale bar, 20 μm. (E) Analysis of percentage of αSMA^+^ cells (N = 30 images per group from 3 HSC cell strains with 3 experimental replicates per HSC cell strain; more than 150 cells were analyzed per cell strain). Symbols with different colors represent different cell strains. The bars and error bars indicate Mean ± SD. Significance was determined by two-way ANOVA using multiple comparisons tests (****p < 0.0001, ns = non-significant).

Collectively, these data demonstrate that TGFβ2 promotes F-actin stress fiber formation in HSC cells, which is decreased by either actin cytoskeleton relaxation or depolymerization. Furthermore, TGFβ2 induces αSMA expression, which is prevented by ROCK inhibition, even as short as 30 min, whereas short-term actin depolymerization does not influence aberrant αSMA stress fiber formation across different HSC cell strains.

### TGFβ2 stabilizes actin cytoskeleton to upregulate YAP/TAZ activity

Recently, we demonstrated that TGFβ2 increases nuclear YAP and TAZ, a readout for active YAP/TAZ signaling, in HTM cells cultured atop or within ECM hydrogels ^26^. To assess the effect of TGFβ2 on YAP/TAZ subcellular localization in HSC cells subjected to the different treatments, we evaluated both YAP and TAZ nuclear-to-cytoplasmic (N/C) ratios. We observed that exposure to TGFβ2 significantly increased YAP N/C ratio (**Fig. 4A,B**), TAZ N/C ratio (**Fig. 4C,D**) and expression of their putative downstream effector TGM2 (**Fig. 4E,F**) compared to controls, suggesting that TGFβ2 enhanced YAP/TAZ transcriptional activity. Given that actin cytoskeletal integrity is required for proper YAP/TAZ regulation in a variety of cells ^41,49^, we investigated whether TGFβ2-induced YAP/TAZ activation in HSC cells depended on an intact actin cytoskeleton. Our results revealed that HSC cells co-treated with Y27632 or Lat B significantly decreased TGFβ2-induced YAP/TAZ nuclear localization and TGM2 expression. Consistent with our observation on F-actin fibers in response to the different treatments, Lat B was more effective in decreasing YAP/TAZ activity in HSC cells (i.e., below control levels) compared to Y27632 (**Fig. 4A-F**).

**Fig. 4.**
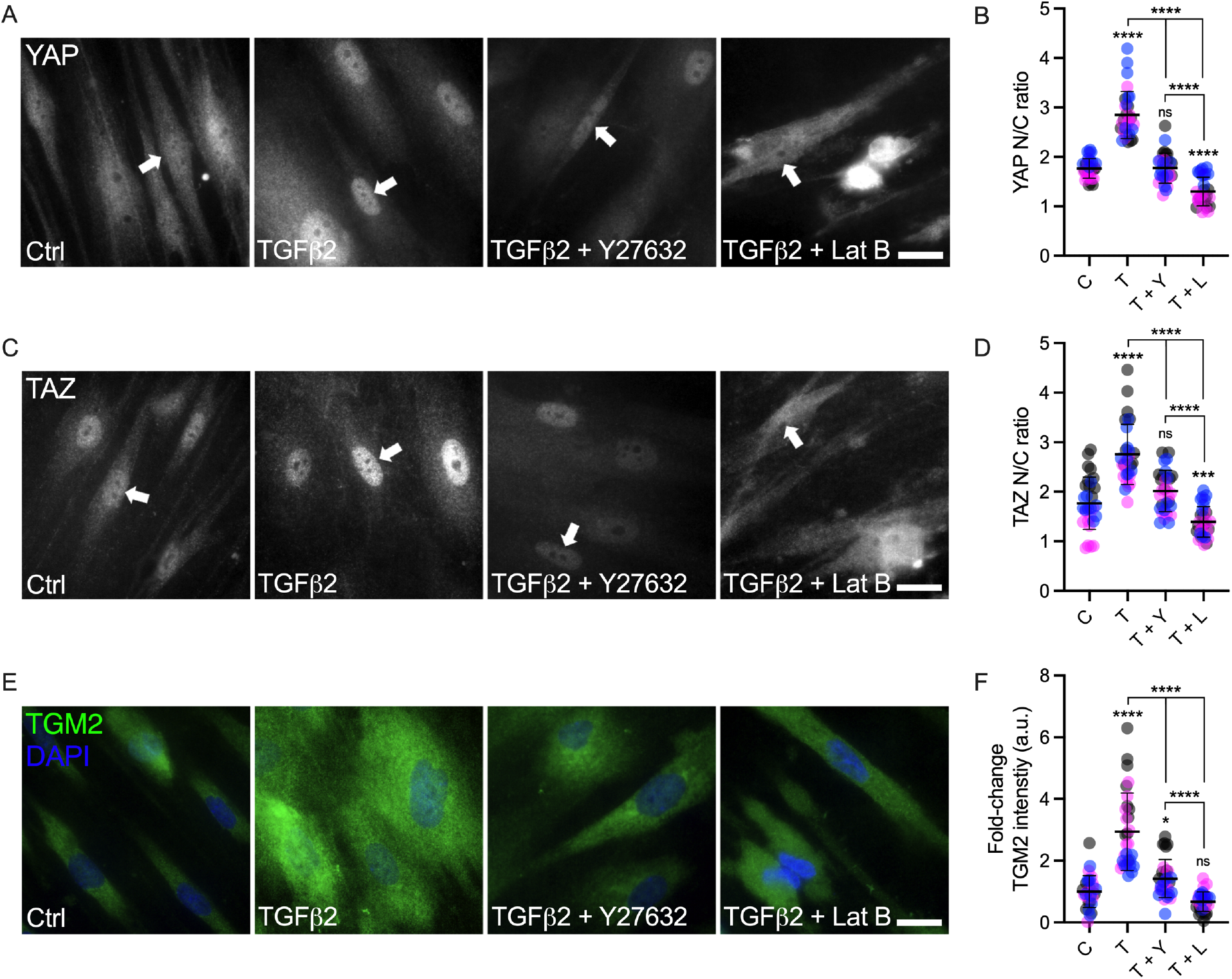
Effects of TGFβ2 in the absence or presence of a ROCK inhibitor or Lat B on YAP/TAZ activity in HSC cells. (A and B) Representative immunofluorescence micrographs of YAP and TAZ in HSC cells on ECM hydrogel substrates subjected to control, TGFβ2 (3 d; 2.5 ng/mL), TGFβ2 + Y27632 (3 d; 10 μM), TGFβ2 (3 d) + Lat B (30 min; 2 μM). Arrows indicate YAP/TAZ nuclear localization. Scale bar, 20 μm. (C and D) Analysis of YAP/TAZ nuclear/cytoplasmic ratio (N = 30 images from 3 HSC cell strains with 3 experimental replicates per cell strain). (E) Representative fluorescence micrographs of TGM2 in HSC cells on ECM hydrogel substrates subjected to the different treatments. Scale bar, 20 μm. (F) Analysis of TGM2 intensity (N = 30 images from 3 HSC cell strains with 3 replicates per cell strain). Symbols with different colors represent different cell strains. The bars and error bars indicate Mean ± SD. Significance was determined by two-way ANOVA using multiple comparisons tests (*p<0.05, ****p < 0.0001, ns = non-significant).

Together, these data show that TGFβ2 increases nuclear YAP/TAZ and TGM2 expression in HSC cells from multiple donors, which is potently attenuated by either actin cytoskeleton relaxation or depolymerization.

### YAP/TAZ mediate ECM remodeling and actomyosin cell cytoskeleton

It has been shown that HSC cells isolated from glaucomatous eyes exhibit increased F-actin stress fibers and expression of ECM proteins including fibronectin ^15^; in this study, our results suggest that TGFβ2 also increases actin cytoskeleton remodeling. Both abnormal ECM deposition and actin remodeling may be part of a pathologic signature in HSC cells. Therefore, we next tested whether pharmacologic YAP/TAZ inhibition could rescue TGFβ2-induced HSC cell dysfunction. To do so, HSC cells atop ECM hydrogels were treated with TGFβ2 alone or co-treated with verteporfin (without light stimulation), which disrupts nuclear YAP/TAZ-TEAD interactions thereby inhibiting transcriptional activity ^50^. Co-treatment with TGFβ2 + verteporfin significantly decreased N/C ratios of YAP and TAZ in HSC cells compared to TGFβ2 alone, approximating baseline levels, in agreement with our recent data using HTM cells ^51^ (**Suppl. Fig. 1A-D**). Similarly, verteporfin significantly decreased TGFβ2-stimulated TGM2 expression; yet, levels remained significantly higher compared to controls (**Suppl. Fig. 1E,F**).

We observed that exposure to verteporfin significantly reduced TGFβ2-induced fibronectin deposition approximating untreated controls (**Fig. 5A,B**). Importantly, we demonstrated that TGFβ2-induced αSMA expression (**Fig. 5C,D**), F-actin stress fiber formation (**Fig. 5E,F**), and phospho-myosin light chain (p-MLC) levels (**Fig. 5G,H**) were significantly decreased by verteporfin co-treatment, but again did not reach baseline levels.

**Fig. 5.**
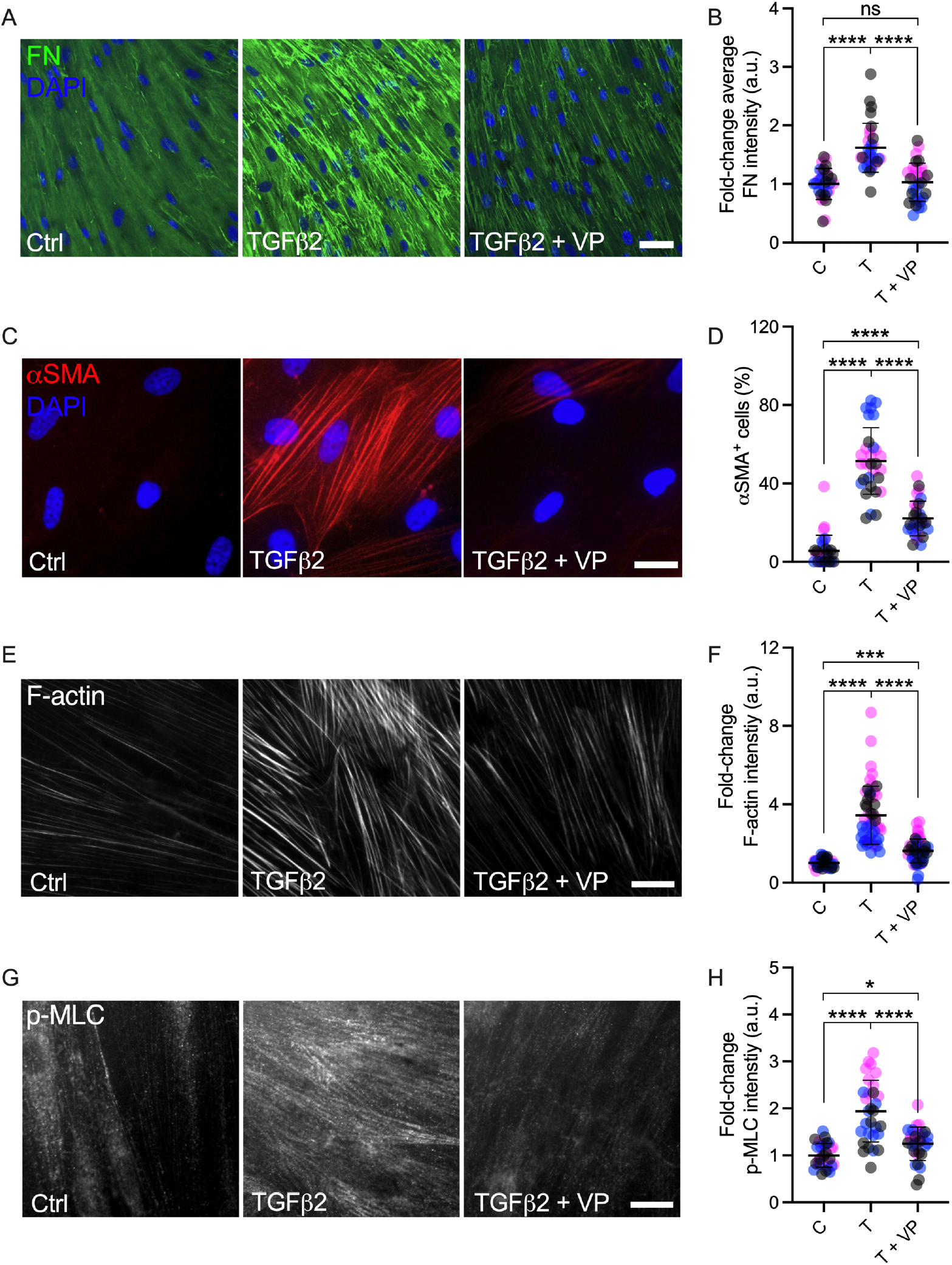
Effects of pharmacologic YAP/TAZ inhibition on ECM remodeling and actomyosin cytoskeleton in HSC cells. (A) Representative immunofluorescence micrographs of fibronectin (FN) in HSC cells on ECM hydrogel substrates subjected to control, TGFβ2 (3 d; 2.5 ng/mL), TGFβ2 + verteporfin (3 d; 0.5 μM). Scale bar, 100 μm. (B) Analysis of fibronectin intensity (N = 30 images per group from 3 HSC cell strains with 3 experimental replicates per HSC cell strain). (C) Representative immunofluorescence micrographs of αSMA in HSC cells on ECM hydrogel substrates subjected to the different treatments. Scale bar, 20 μm. (D) Analysis of percentage of αSMA^+^ cells (N = 3 HSC cells strains, more than 150 cells were analyzed per cell strain). (E and G) Representative fluorescence micrographs of F-actin and p-MLC in HSC cells on ECM hydrogel substrates subjected to the different treatments. Scale bar, 20 μm. (F and H) Analysis of F-actin and p-MLC intensity (N = 30 images from 3 HSC cell strains with 3 experimental replicates per cell strain). Symbols with different colors represent different cell strains. The bars and error bars indicate Mean ± SD. Significance was determined by two-way ANOVA using multiple comparisons tests (*p<0.05, ***p<0.001, ****p < 0.0001, ns = non-significant).

In sum, these data suggest that pharmacologic YAP/TAZ inhibition reduces the expression of fibronectin and decreases actomyosin cytoskeletal rearrangement across different HSC cell strains.

### YAP/TAZ mediate TGFβ2-induced HSC cell contractility

It has been shown that perfusing *ex vivo* human anterior segments with TGFβ2 reduced the length of the SC inner wall ^27^, suggestive of tissue contraction. HSC cells are known to be highly contractile ^3^. In support of this notion, we observed that HSC cell-laden hydrogels - an experimental tool developed in our laboratory to characterize HTM cell behavior ^26,30,43^ - in absence of any treatment significantly contracted over time, reaching ~35-42% of their original size by 5 d with normal donor-to-donor variability (**Suppl. Fig.2**).

Our data so far showed that YAP/TAZ inhibition reduced pathologic actomyosin cytoskeleton remodeling. Therefore, we hypothesized that TGFβ2 may increase HSC cell contractility, which could be reduced by YAP/TAZ inhibition. To test this hypothesis, we again encapsulated HSC cells in ECM hydrogels and treated the constructs with TGFβ2, either alone or in combination with verteporfin (without light stimulation), to assess the level of hydrogel contraction at 5 d. TGFβ2-treated HSC hydrogels exhibited significantly greater contraction compared to controls (62.79% of controls; **Fig. 6A,B**), consistent with our previous studies using HTM cells ^26,30,43^. Co-treatment with verteporfin significantly decreased pathologic HSC hydrogel contraction (81.56% of controls) compared to TGFβ2-treated samples (**Fig. 6A,B**), but it did not fully restore baseline levels. Of note, the influence of verteporfin on HSC cell-laden hydrogel contraction differed between HSC cell strains, showing a stronger effect on HSC02 than HSC03 and HSC09 in line with normal donor-to-donor variability (**Fig. 5A,B**). To rule out that hydrogel contractility was influenced by the cell number, we assessed HSC cell viability in constructs subjected to the different treatments. No differences were observed for HSC cell-laden hydrogels across groups (**Fig. 6C**).

**Fig. 6.**
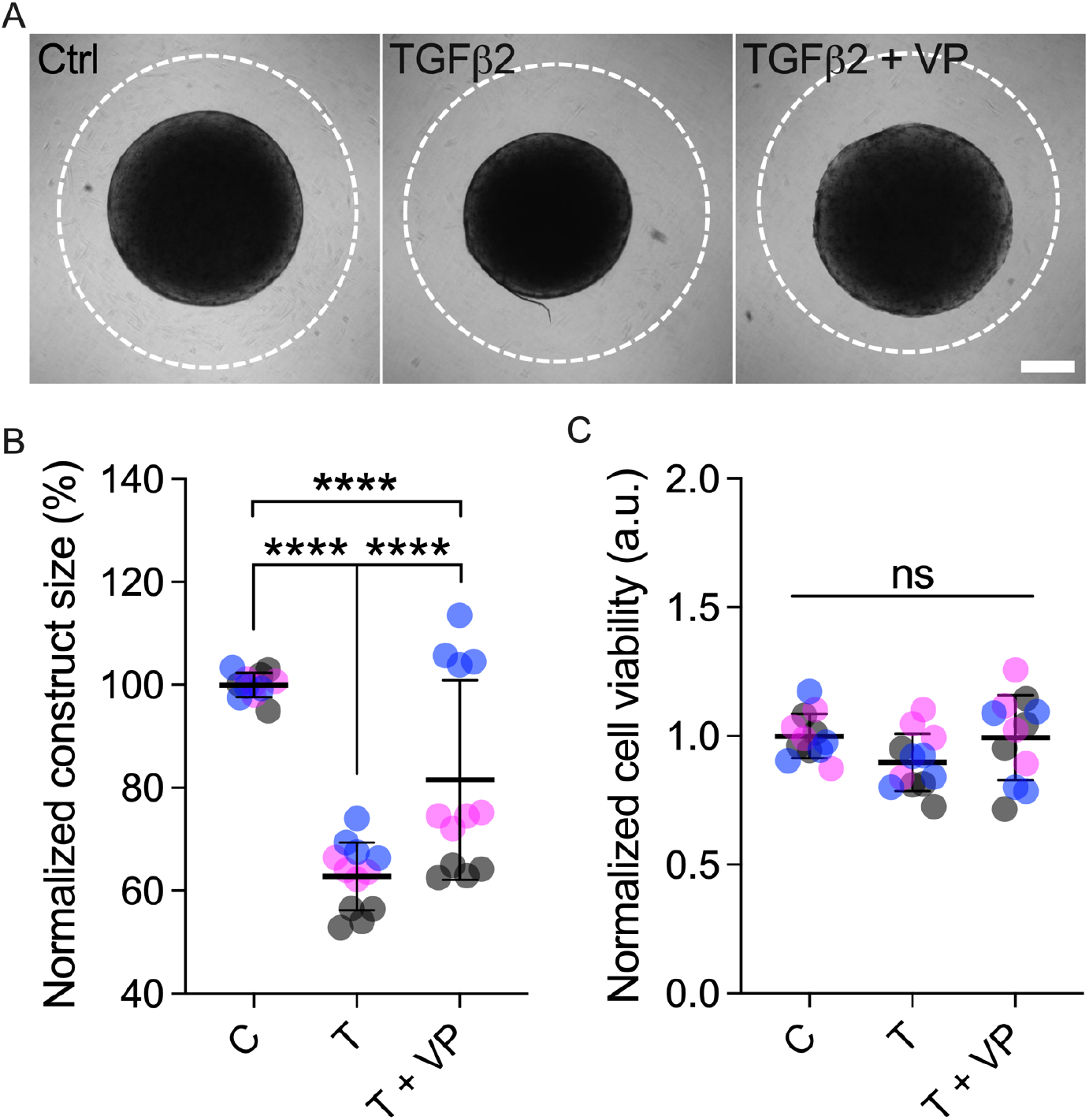
Effects of TGFβ2 in the absence or presence of YAP/TAZ inhibition on HSC hydrogel contractility. (A) Representative brightfield images of HSC hydrogels subjected to control, TGFβ2 (3 d; 2.5 ng/mL), TGFβ2 + verteporfin (3 d; 0.5 μM) at 5 d. Dashed lines outline original size of constructs at 0 d. Scale bar, 1 mm. (B) Construct size quantification of HSC hydrogels subjected to the different treatments at 5 d (N = 12 hydrogels per group from 3 HSC cell strains with 4 experimental replicates per HSC cell strain). (C) Cell viability quantification of HSC hydrogels subjected to the different treatments at 5 d (N = 12 hydrogels per group from 3 HSC cell strains with 4 experimental replicates per HTM cell strain). Symbols with different colors represent different cell strains. The bars and error bars indicate Mean ± SD. Significance was determined by two-way ANOVA using multiple comparisons tests (****p<0.0001, ns = non-significant).

Together, these data demonstrate that TGFβ2 robustly induces HSC cell contractility in a 3D ECM microenvironment across multiple cell strains, and that pharmacologic YAP/TAZ inhibition potently decreases pathologic HSC cell contraction.

## Discussion

The profibrotic cytokine TGFβ2 is a key contributor to outflow tissue dysfunction in POAG ^10,19^. TGFβ2 has been shown to increase the deposition of ECM material in the JCT-TM beneath the SC inner wall endothelium and contract the SC ^26,27,30^. Most *in vitro* studies to date investigating the effects of TGFβ2 on outflow cell dysfunction have focused on TM cells ^26,30,48^. In contrast, the contributions of TGFβ2 to SC cell pathobiology are considerably less well understood. A recent multi-ethnic genome wide meta-analysis identified *YAP1* as a potential genetic risk factor for POAG, implicating that YAP (and perhaps TAZ by association) may play a critical role in glaucoma pathogenesis ^40^. We have shown that TGFβ2 upregulates YAP/TAZ activity in HTM cells cultured atop or within ECM hydrogels ^26^. Here, our aim was to elucidate mechanisms governing YAP/TAZ modulation in HSC cells in response to TGFβ2 by culturing cells on a tissue-mimetic ECM substrate.

Glaucomatous SC cells isolated from POAG eyes have been shown to exhibit more F-actin stress fibers with an overall higher degree of disorganization compared to normal cells ^15^. Consistent with this observation, we found that TGFβ2 significantly increased F-actin stress fiber formation in HSC cells with qualitatively more disorganized filaments (**Fig. 2B,C**). Concurrent actin destabilization with Y27632 or Lat B potently decreased TGFβ2-induced actin stress fibers, with short-term Lat B exposure (30 min; necessary to preserve HSC cell viability) showing a stronger effect compared to ROCK inhibitor co-treatment for the full 3 days. When Y27632 was supplemented for only 30 min (i.e., matching the Lat B regimen) to allow for a more accurate side-by-side comparison of the two compounds, an even greater inhibitory effect on pathologic F-actin stress fiber formation was observed (30 min: ~80.94%; 3 d: ~49.55% vs. TGFβ2; **Fig. 2B,C** and **Fig. 3B,C**). These data raise the possibility that by day 3, Y27632 may have lost some of its potency; this would be in line with the reported Y27632 half-life of ~12-16 h ^52^. Nevertheless, it is important to note that despite potentially weakened ROCK inhibitory activity, co-treatment of TGFβ2 + Y27632 significantly decreased pathologic F-actin disorganization in HSC cells. It has been extensively reported that the glaucoma-associated stressors dexamethasone and TGFβ2 stimulate the formation of cross-linked actin networks (CLANs) in HTM cells ^53–56^. In this study, we did not observe any such structures in HSC cells independent of the treatment applied. A number of signaling pathways including Smad, Wnt, ROCK and ERK have been shown to modulate CLAN formation in HTM cells ^57^. Further research will be necessary to investigate the specific contribution of these important signaling pathways in HSC cell pathobiology in the context of cytoskeletal homeostasis and remodeling.

The transcriptional coactivators YAP and TAZ play important roles in mechanotransduction and POAG pathogenesis ^40,41^. Recently, we showed that activity of YAP/TAZ is upregulated in HTM cells under diverse simulated glaucomatous conditions ^26^. However, the mechanisms underlying YAP/TAZ signaling in HSC cells influenced by TGFβ2 remain to be elucidated. We here demonstrated that TGFβ2 increased YAP/TAZ nuclear localization and activity in HSC cells, which was potently attenuated by actin cytoskeleton relaxation using Y27632 or actin depolymerization using Lat B (**Fig. 4**). This suggests that actin integrity is required for YAP/TAZ activation in HSC cells. We found that Lat B treatment was more effective in blocking TGFβ2-induced YAP/TAZ nuclear accumulation (i.e., more cytoplasmic YAP/TAZ) than ROCK inhibition with Y27632, consistent with the F-actin stress fiber data. A previous study showed that HTM cells can recover to their initial state after removal of Lat B treatment in a short period of time ^58^. It would be worthwhile to further investigate whether Y27632/Lat B-reduced YAP/TAZ activity could be reversed with the treatments withheld.

Endothelial-to-mesenchymal transition (EndMT) is a process whereby endothelial cells undergo a series of molecular events, such as increased expression of αSMA, fibronectin, vimentin, and collagen types I and III, that lead to a change in phenotype toward mesenchymal-like cells ^59,60^. TGFβ has been shown to induce EndMT, which can contribute to various fibrotic diseases and cancers ^60–63^. In a previous study, it was shown that TGFβ2 stimulated EndMT of HSC cells cultured on top of a synthetic biomaterial; downregulation of endothelial cell markers and upregulation of mesenchymal makers were noted ^64^. Consistent with this, we showed that TGFβ2 enhanced EndMT of HSC cells cultured atop tissue-mimetic ECM hydrogels, as indicated by increased formation of fibronectin and αSMA (**Fig. 2D,E, Fig. 3D,E, Fig. 5A-D**). The TGFβ2-induced αSMA stress fibers were potently disrupted by short-term exposure to Y27632. We measured a ~71.84% decrease in αSMA-positive cells with only 30-min of ROCK inhibition compared to TGFβ2-treated HSC cells, nearly identical to the full 3 d co-treatment (~69.52%). By contrast, Lat B-mediated actin depolymerization failed to reduce αSMA-positive cells. Importantly, we found that TGFβ2-induced EndMT in HSC cells was potently blocked by pharmacologic YAP/TAZ inhibition with verteporfin in absence of light stimulation consistent with previous reports in non-ocular ^65–67^ and ocular cells ^51,68,69^; significantly decreased fibronectin and actin stress fibers were noted (**Fig. 5A-D**). Moreover, TAP/TAZ inhibition decreased TGFβ2-induced expression of TGM2 (**Suppl. Fig. 1E,F**), which crosslinks fibronectin and thereby stiffens the ECM ^70^. This suggests that YAP/TAZ inhibition may decrease ECM stiffness to improve aqueous humor outflow. Therefore, targeting YAP/TAZ signaling in the SC endothelium to inhibit EndMT is an intriguing strategy for managing ocular hypertension.

Actomyosin, the actin-myosin complex, regulates cell contractility in various cell types ^71^ including HSC cells ^3^. To that end, HSC cells in absence of any treatment exhibited qualitatively fewer F-actin fibers compared to HTM cells when cultured atop our ECM hydrogels. Therefore, we speculate that the higher degree of HSC cell contractility could stem from upregulated activity of myosin II, which is responsible for producing contraction force ^72^. Myosin II activity is primarily regulated by phosphorylation of MLC; this process is mediated by myosin light chain kinase (MLCK) and myosin phosphatase ^73^. Importantly, MLCK inhibition has been shown to decrease IOP in rabbit eyes ^74^. Blebbistatin, a pharmacologic inhibitor of myosin II adenosine triphosphatase activity, increases outflow facility via blocking the binding of myosin to actin ^75^. Futures studies would be necessary to investigate in greater detail the role of myosin II activity and myosin-actin interactions in HSC cell biology.

Consistent with our observation on HTM cells, TGFβ2-treated HSC cells exhibited elevated levels of cell contractility-related molecules (i.e., αSMA, F-actin and p-MLC), which correlated with increased HSC hydrogel contraction. Importantly, this TGFβ2-induced pathologic process was potently attenuated by pharmacologic YAP/TAZ inhibition (**Fig. 5C-H, Fig. 6B-D**). Previous studies showed that outflow resistance is modulated by SC cell stiffness, which is directly correlated with their contractility status ^14,76^. It would be worthwhile to further investigate the effects of YAP/TAZ inhibition on HSC cell stiffness.

In conclusion, by culturing HSC cells on tissue-mimetic ECM hydrogels, we demonstrated that TGFβ2 drives actin stress fiber formation to upregulate YAP/TAZ activity, and that YAP/TAZ inhibition reduces HSC cell contractility, which may positively affect cell stiffness and outflow resistance. Our findings provide new evidence for a pathologic role of YAP/TAZ hyperactivation in HSC cell pathobiology in glaucoma, and suggest that pharmacologic YAP/TAZ inhibition has promising potential to improve outflow tissue dysfunction.

## Acknowledgments

We thank Dr. Robert W. Weisenthal and the team at Specialty Surgery Center of Central New York for assistance with corneal rim specimens. We also thank Dr. Nasim Annabi at the University of California – Los Angeles for providing the KCTS-ELP, Dr. Alison Patteson at Syracuse University for rheometer access, and Drs. Audrey M. Bernstein and Mariano S. Viapiano at SUNY Upstate Medical University for imaging support.

## Author contributions

H.L., A.S., K.M.P., W.D.S., P.S.G., and S.H. designed all experiments, collected, analyzed, and interpreted the data. K.M.P. and W.D.S. provided the HSC reference cells. H.L. and S.H. wrote the manuscript. All authors commented on and approved the final manuscript. P.S.G. and S.H. conceived and supervised the research.

## Data and materials availability

All data needed to evaluate the conclusions in the paper are present in the paper and/or the Supplementary Materials. Additional data related to this paper may be requested from the authors.

