## Supplementary information for "YAP/TAZ mediate TGFβ2-induced Schlemm’s canal cell dysfunction"

### *Supplementary Material*

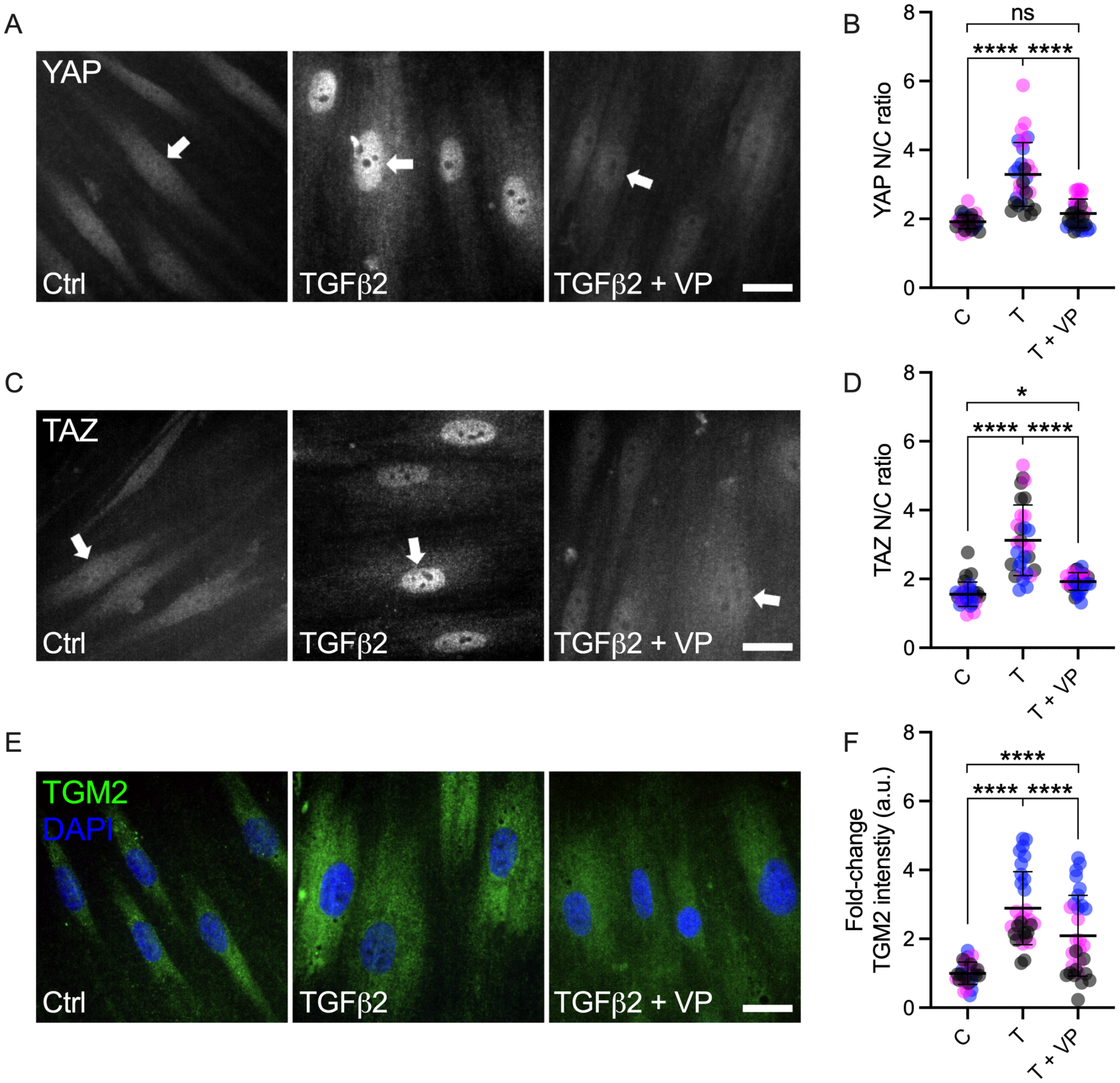


**Suppl. Fig. 1. Effects of TGFβ2 in absence or presence of VP on YAP/TAZ activity in HSC cells.** (A and B) Representative fluorescence micrographs of YAP and TAZ in HSC cells on ECM hydrogel substrates subjected to control, TGFβ2 (3 d; 2.5 ng/mL), TGFβ2 + verteporfin (3 d; 0.5 μM) (YAP/TAZ = grey). Scale bar, 20 μm; arrows indicate YAP/TAZ nuclear localization. (C and D) Analysis of YAP/TAZ nuclear/cytoplasmic ratio (N = 30 images from 3 HSC cell strains with 3 replicates per cell strain). (E) Representative fluorescence micrographs of TGM2 in HSC cells on ECM hydrogel substrates subjected to the different treatments (TGM2 = green; DAPI = blue). Scale bar, 20 μm. (F) Analysis of TGM2 intensity (N = 30 images from 3 HSC cell strains with 3 experimental replicates per cell strain). Symbols with different colors represent different cell strains. The bars and error bars indicate Mean ± SD. Significance was determined by two-way ANOVA using multiple comparisons tests (*<005, ****p < 0.0001, ns = non-significant).

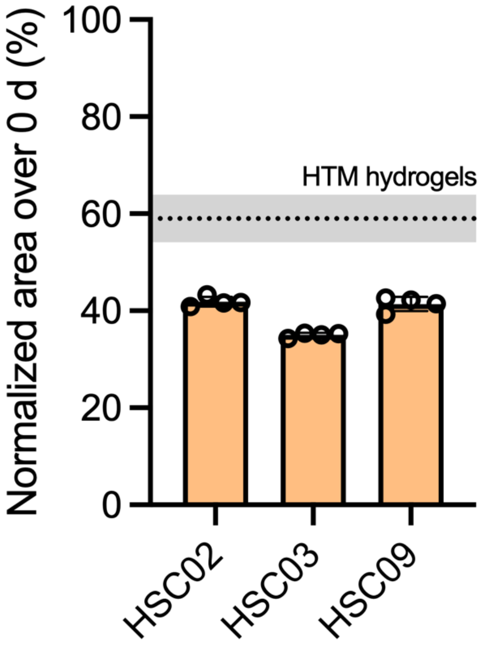


**Suppl. Fig. 2. HSC hydrogel contraction**. Quantification of HSC hydrogel contraction at 5 d (N = 4 experimental replicates per cell strain; dotted line with gray shading = Mean ± SD of HTM hydrogels from refs [26,43]).

**Supplementary Table 1: Antibody information.**

| **Target** | **Catalog no.** | **Company** | **Dilution IB** | **Dilution ICC** |
|---|---|---|---|---|
| anti-Fibulin-2 | ab251662 | Abcam | | 1:2,000 |
| anti-VE-Cadherin | 2500 | Cell Signaling Technology | 1:500 | |
| anti-GAPDH | G9545 | Sigma | 1:80,000 or 1:160,000 | |
| anti-MYOC | MABN866 | Sigma | 1:2,000 | |
| anti-YAP | 14074S | Cell Signaling Technology | | 1:200 |
| anti-TAZ | 4883S | Cell Signaling Technology | | 1:200 |
| anti-Fibronectin | ab45688 | Abcam | | 1:500 |
| Cy3-anti-αSMA | C6198 | Sigma | | 1:400 |
| anti-p-MLC | 3675 | Cell Signaling Technology | | 1:200 |
| anti-TGM2 | ab421 | Abcam | | 1:400 |
| Alexa Fluor® 488-conjugated anti-Rabbit IgG | A27034 | Invitrogen | | 1:500 |
| Alexa Fluor® 584-conjugated anti-Rabbit IgG | ab150080 | Abcam | | 1:500 |
| Alexa Fluor® 488-conjugated anti-Mouse IgG | A21203 | Invitrogen | | 1:500 |
| HRP-conjugated anti-Rabbit IgG | ab6721 | Abcam | 1:20,000 | |
| IRDye® 800CW Goat anti-Mouse IgG | 926-32210 | LI-COR | 1:15,000 | |
